# Kryptix-1: Conformational Ensemble Sampling Substantially Improves Cryptic Pocket Detection Relative to Static-Structure and Prior Computational Approaches

**DOI:** 10.64898/2026.09.16.752177

**Authors:** Ryan Varghese, Pooja Tiwary, Krishil Oswal

**Affiliations:** Kryptix-1 Development Team; Department of Pharmaceutical Sciences, Philadelphia College of Pharmacy, Saint Joseph’s University, Philadelphia, PA, 19104, USA; Covenant Biosciences, 447 Sutter St, San Francisco, CA, 94108, USA

## Abstract

Cryptic binding pockets; sites absent or occluded in a protein’s resting-state structure that become druggable only in specific, transiently populated conformations; represent one of the largest untapped opportunities in structure-based drug discovery. Large-scale structural surveys estimate that cryptic pockets occur in on the order of one in six protein families genome-wide, and that accounting for them could expand the druggable fraction of the disease-associated human proteome from roughly 40% to nearly 80%. Detecting cryptic pockets computationally has historically forced a choice between physics-based conformational sampling, which is reliable but too computationally expensive to deploy across more than a handful of targets, and fast machine-learning pocket predictors trained on static structures, which frequently over-predict and lose the precision needed for practical triage. We report Kryptix-1, an in-house pipeline developed at Covenant Biosciences that combines conformational ensemble generation with a consensus pocket-scoring algorithm, benchmarked here against a matched static-structure baseline and against the published literature on a 10-protein sample from CryptoBench, an independently curated cryptic-pocket reference dataset. On the field’s own standard residue-overlap metric (Jaccard index ≥ 0.5), Kryptix-1 succeeds on 80% of benchmark proteins, compared to 20% for the static baseline and approximately 40–45% for the best-performing methods reported in an independent comparative evaluation using the same metric. We report these results alongside an explicit accounting of where Kryptix-1’s advantage is smaller or reverses, and we are explicit throughout that this is a small pilot evaluation, not a fully powered validation study.

## Introduction

The central limiting factor in modern structure-based drug discovery is not chemistry but geometry: a compound cannot be designed against a binding site that is not there to see. The concept of the “druggable genome,” the fraction of gene products with structural features amenable to small-molecule binding, was formalized over two decades ago and estimated to cover only a minority of disease-relevant human proteins [1]. By conservative modern estimates, on the order of 80–85% of proteins encoded by the human genome present no obvious, well-formed pocket in their resting-state structure and remain conventionally classified as undruggable [2]. This has direct, quantifiable downstream consequences. Target selection failures are implicated in a substantial fraction of late-stage clinical attrition [3, 4]. Once attrition across the full pipeline is accounted for, the aggregate cost of bringing a single new molecular entity to market has been estimated at $1.8 billion [4]. A meaningful share of this attrition is not a failure of chemistry against a validated, well-characterized site; it is the absence of a validated, well-characterized site to begin with.

The “undruggable” classification, however, is a statement about static structures, not about proteins as physical objects. A substantial and growing body of evidence shows that many apparently pocket-less proteins undergo transient conformational excursions that open genuine, druggable cavities, cryptic pockets, which simply do not appear in a crystal structure, a cryo-EM map, or a single AlphaFold prediction [5, 6]. The 2016 CryptoSite study demonstrated that accounting for this class of site alone could expand the druggable fraction of the disease-associated human proteome from roughly 40% to as much as 78% [7], very nearly doubling the addressable target space without a single new chemical modality. More recent large-scale structural surveys have placed the prevalence of cryptic pockets at approximately 16% of protein clusters genome-wide, with a meaningful fraction of currently “undruggable” proteins harboring a buried, ligandable, and in some cases fully druggable cryptic site [8]. Beyond simply expanding addressable surface area, cryptic and allosteric pockets have already yielded validated pharmacological starting points: engineered small molecules targeting a cryptic allosteric site have been shown to produce both positive and negative modulation of target activity in a real protein system [9], and systematic structural analysis across dozens of proteins with experimentally confirmed cryptic sites has begun to distinguish genuinely cryptic pockets from simple allosteric coupling, a distinction with direct consequences for how a given site should be pursued therapeutically [10].

The obstacle has never been that cryptic pockets are rare. It is that finding them is hard, and finding them at the scale a real discovery program requires has been harder still. The physical origin of a cryptic pocket is a rare, often high-energy conformational state; side-chain repacking, partial secondary-structure unfolding, or loop rearrangement; that a protein visits only a small fraction of the time [5]. Mapping these states computationally has conventionally required long-timescale molecular dynamics or enhanced sampling, frequently on the order of hundreds of microseconds of aggregate simulation time per target, since the slow motions responsible for cryptic pocket opening routinely fall on microsecond-to-millisecond timescales well beyond conventional simulation protocols [5, 11]. Enhanced-sampling variants, including replica-exchange methods with scaled solvent-protein interactions, have improved the efficiency of this search considerably [12], but the underlying computational burden remains substantial enough that physics-based cryptic pocket discovery has largely stayed confined to single-target, hypothesis-driven case studies rather than multi-target screening campaigns.

The natural response, and the dominant trend in the field over the past several years, has been to replace expensive simulation with fast, learned predictors. This landscape has evolved considerably over roughly fifteen years: early geometric cavity-detection algorithms based on alpha-sphere clustering [13] gave way to machine-learning re-ranking approaches trained on labeled structural features [14], and more recently to graph neural networks trained end-to-end to flag cryptic-pocket-forming residues directly from a single input structure, without any conformational sampling at all [15]. This trajectory has been accelerated further by using AlphaFold-predicted structures as a substitute for experimental starting points [6, 16]. These methods are fast: inference on a single structure can take seconds rather than the hundreds of GPU-hours a full ensemble simulation requires. But they inherit a structural limitation from their input. AlphaFold and comparable structure predictors are trained to output a single, most-probable conformation, and are known to struggle to represent the alternative, functionally important states that many proteins genuinely populate [11, 16]. A recent, large-scale comparative evaluation spanning thirteen binding-site predictors and thirty years of methodological development found that even the best-performing current methods top out at a residue-level F1 score of approximately 0.42, with classical geometric approaches scoring as low as 0.23 [17]. The practical consequence, familiar to anyone who has triaged the output of these tools, is a precision problem: methods tuned to maximize recall on this genuinely hard task tend to flag a large fraction of a protein’s surface as candidate pocket, producing lists that are directionally useful but require extensive manual curation before they are actionable. Consensus approaches, combining the output of multiple independent detection or scoring methods rather than relying on any single one, have shown consistent, reproducible improvements over single-method predictions in the closely related problem of docking pose and virtual screening enrichment [18], motivating a similar strategy for pocket detection specifically.

Kryptix-1 was built to resolve this tension directly, rather than to trade one side of it for the other. It retains the physical grounding of conformational sampling; the recognition that a cryptic pocket is fundamentally a dynamical, not a static, property of a protein; while remaining fast enough to deploy at the scale a real screening program requires, running on a single consumer-grade GPU for the benchmarking reported here. This report benchmarks Kryptix-1 against a matched static-structure baseline (identical scoring logic, no conformational sampling) on a randomly selected 10-protein sample drawn from CryptoBench [19], a published, independently curated dataset of cryptic protein-ligand binding sites, and situates the results against previously reported findings in the independent literature. We report three metrics adopted directly from established conventions in the pocket-prediction field; Jaccard residue overlap, distance-to-correct-center, and residue-level precision/recall; rather than an evaluation framework of our own design, so that the results can be read directly against prior work. We are explicit throughout this report that the evaluation described here is a pilot: ten proteins, one benchmark dataset, and a single consumer-grade GPU, evaluated without the statistical power a fully confirmatory study would require. This report does not include three specific validations:

- full validation against CryptoBench’s complete published collection of over 1,100 structures;
- wet-laboratory confirmation of any individual predicted site; and
- benchmarking on dedicated enterprise-scale compute infrastructure.

What follows is intended as an early, honestly-scoped signal that reintroducing conformational dynamics into cryptic pocket detection is a tractable and substantially rewarding direction, not as a final validation of Kryptix-1 as a deployed discovery tool.

## Results

### Jaccard residue overlap

At the field-standard success threshold of Irel ≥ 0.5, Kryptix-1 succeeded on 8 of the 10 benchmark proteins (80%), against 2 of 10 (20%) for the static-structure baseline (Table 1, Fig. 1). This difference was assessed with an exact McNemar test on the paired per-protein success/failure outcomes (6 discordant pairs, all in Kryptix-1’s favor, in the baseline’s favor; p = 0.031), and is significant at the conventional 0.05 threshold despite the small sample. This compares favorably to the 40–45% success rate reported for the best-performing methods in an independent large-scale comparative evaluation using the same threshold convention [17].

**Table 1.** Jaccard Overlap Success Rate (Irel 0.5). Exact McNemar test, Kryptix-1 vs. static baseline, paired by protein (n = 10).

| Approach | Successes | Success Rate | p-value |
| --- | --- | --- | --- |
| Static-structure baseline | 2 / 10 | 20% | N/A |
| Kryptix-1 | 8 / 10 | 80% | 0.031 (see note) |
| Best published methods [17] | N/A | 40–45% | N/A |

**Figure 1.**
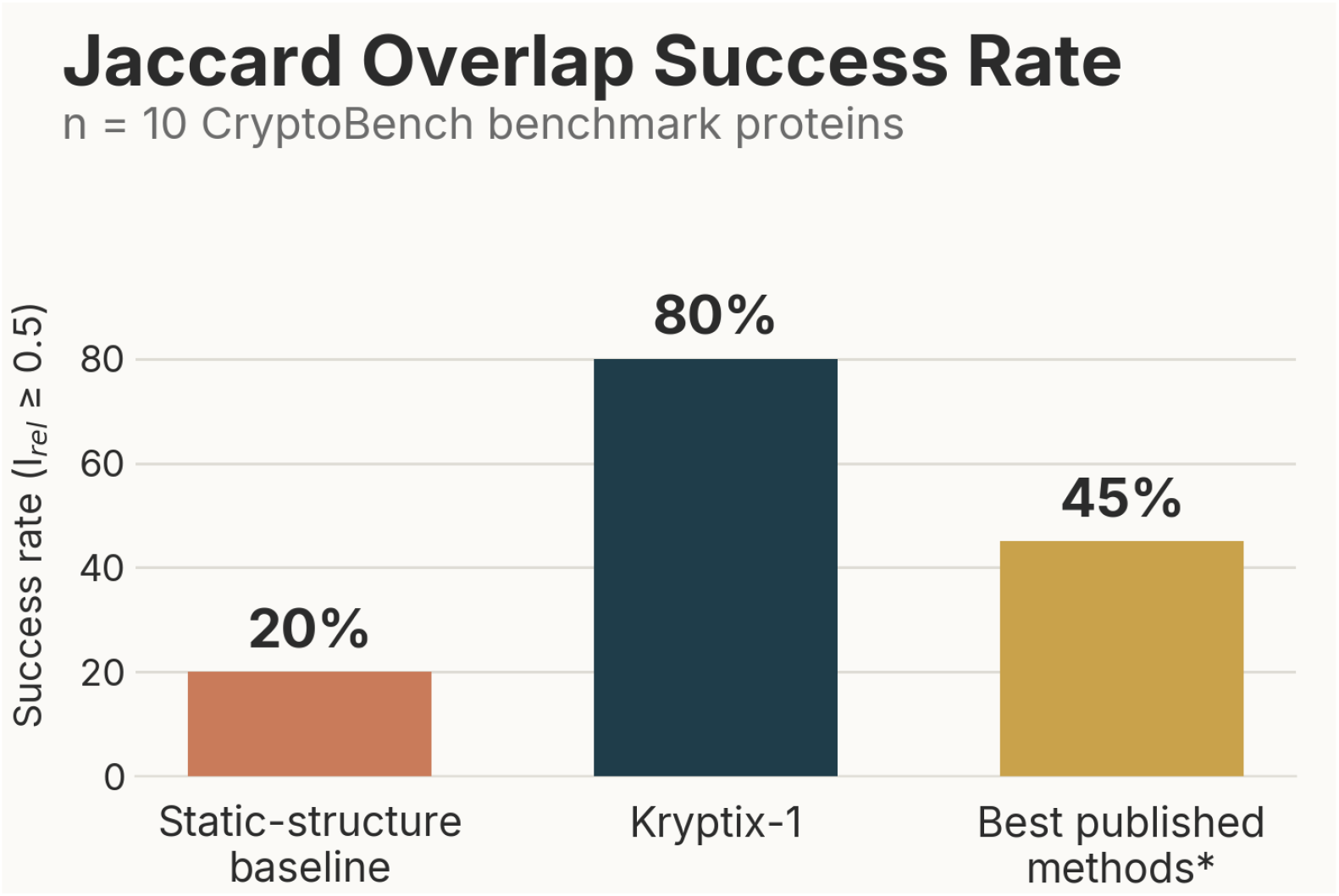
Jaccard overlap (Irel ≥0.5) success rate across the 10-protein benchmark sample, for the static-structure baseline, Kryptix-1, and the best-performing methods reported in an independent comparative evaluation using the same threshold convention [17].

### Distance to correct center (DCC)

At the strict DCC ≤ 4 Å threshold, Kryptix-1 succeeded on 8 of 10 proteins (80%) against 5 of 10 (50%) for the static baseline (Table 2, Fig. 3); an exact McNemar test on this same paired outcome (3 discordant pairs, all in Kryptix-1’s favor) gives p = 0.25, which does not reach significance at this sample size; we report this honestly rather than selectively citing only the significant Jaccard result above, and return to the small-sample caveat in the Limitations subsection below. At the looser DCC ≤ 12 Å threshold used in some prior benchmark comparisons, both approaches reached 100% success on this sample, a ceiling effect that is not discriminating at this threshold for this particular benchmark. For context, the best previously published single-method results at this same looser threshold fall in the range of 52–60% [17]; we do not present this specific comparison as a controlled head-to-head, since the published figures were obtained on a different dataset under a different evaluation protocol, and note this explicitly as a limitation below.

**Table 2.** DCC Success Rate at Two Thresholds. Exact McNemar test, Kryptix-1 vs. static baseline, paired by protein (n = 10).

| Approach | DCC $\leq 4$ Å | p-value | DCC $\leq 12$ Å |
| --- | --- | --- | --- |
| Static-structure baseline | 50% | N/A | 100% |
| Kryptix-1 | 80% | 0.25 (see note) | 100% |

Fig. 2 shows the per-protein Jaccard overlap underlying the aggregate result above, and Fig. 3 shows the corresponding DCC success rates.

**Figure 2.**
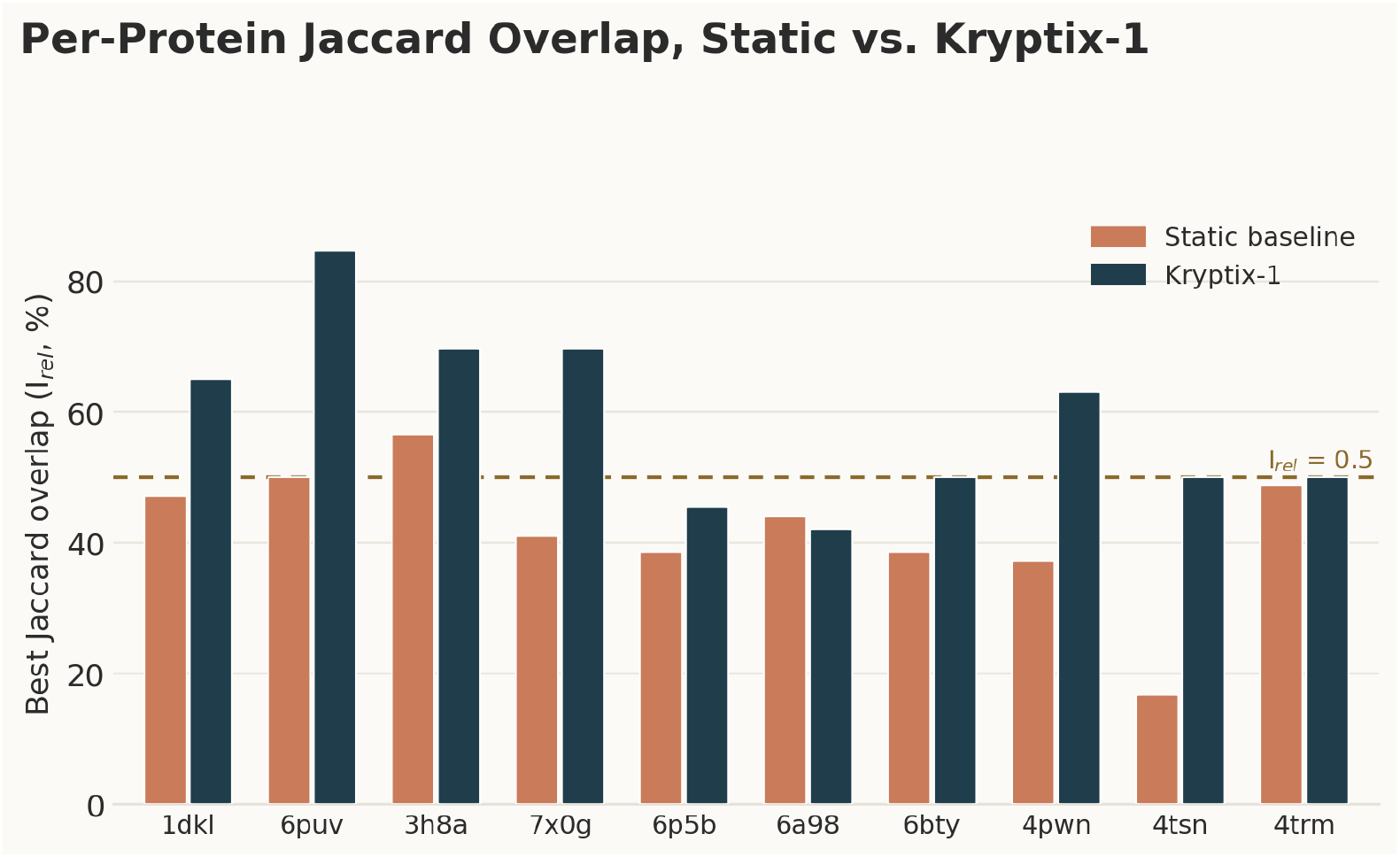
Per-protein best-achieved Jaccard overlap, static baseline versus Kryptix-1, across all 10 benchmark proteins. Kryptix-1 improves on the static baseline for 9 of 10 proteins and crosses the Irel ≥ 0.5 success threshold (dotted line) on 8; one protein (6a98) shows a small, honestly-reported reversal in the static baseline’s favor.

Only two of the ten benchmark proteins failed to reach DCC ≤ 4 Å success under either approach, indicating these represent genuinely difficult cases for both static and ensemble-based detection rather than a systematic weakness specific to either method.

### Recall–precision tradeoff

Kryptix-1 improves mean residue-level recall substantially relative to the static baseline (83.9% → 98.6%, +14.7 percentage points, reaching near-total ground-truth pocket coverage across the benchmark sample; paired t-test t = 2.17, p = 0.058, n = 10; Wilcoxon signed-rank p = 0.063), with a corresponding reduction in mean residue-level precision (28.5% 19.3%, 9.2 percentage points; paired t-test t = −3.68, p = 0.005; Wilcoxon signed-rank p = 0.027; Fig. 4, error bars show ±1 SEM across the 10 proteins). The recall improvement narrowly misses the conventional 0.05 threshold, consistent with a genuine but modest effect measured on a small sample rather than a null result; the static baseline is already close to a recall ceiling on several proteins in this sample, which compresses the paired differences available to the test. We discuss the interpretation of this tradeoff, and why we do not consider it to undermine the two success-rate results above, in the Discussion. For context, an independent large-scale comparative evaluation reports residue-level F1 scores ranging from 0.23 for classical geometric approaches to 0.42 for the best-performing current machine-learning methods on a comparable, non-cryptic binding-site prediction task [17], and CryptoBench’s own baseline evaluation of established cryptic-pocket predictors reports area-under-precision-recall-curve values in the 0.19–0.21 range [19]. Precision and recall in the range reported here, for both approaches, are consistent with this published performance ceiling rather than a departure from it.

**Figure 3.**
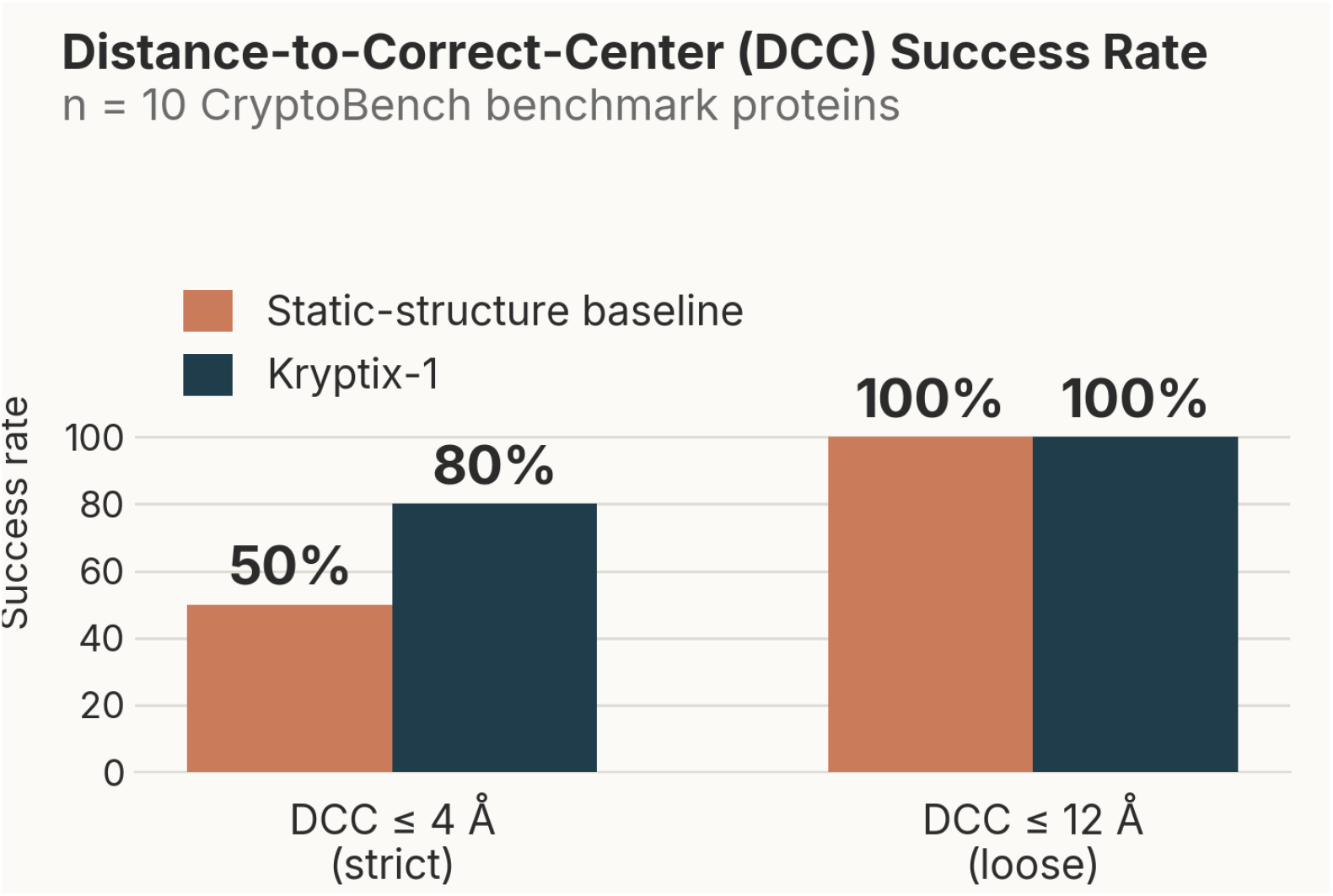
DCC success rate at the strict (4 Å) and loose (12 Å) thresholds, static
baseline versus Kryptix-1.

**Figure 4.**
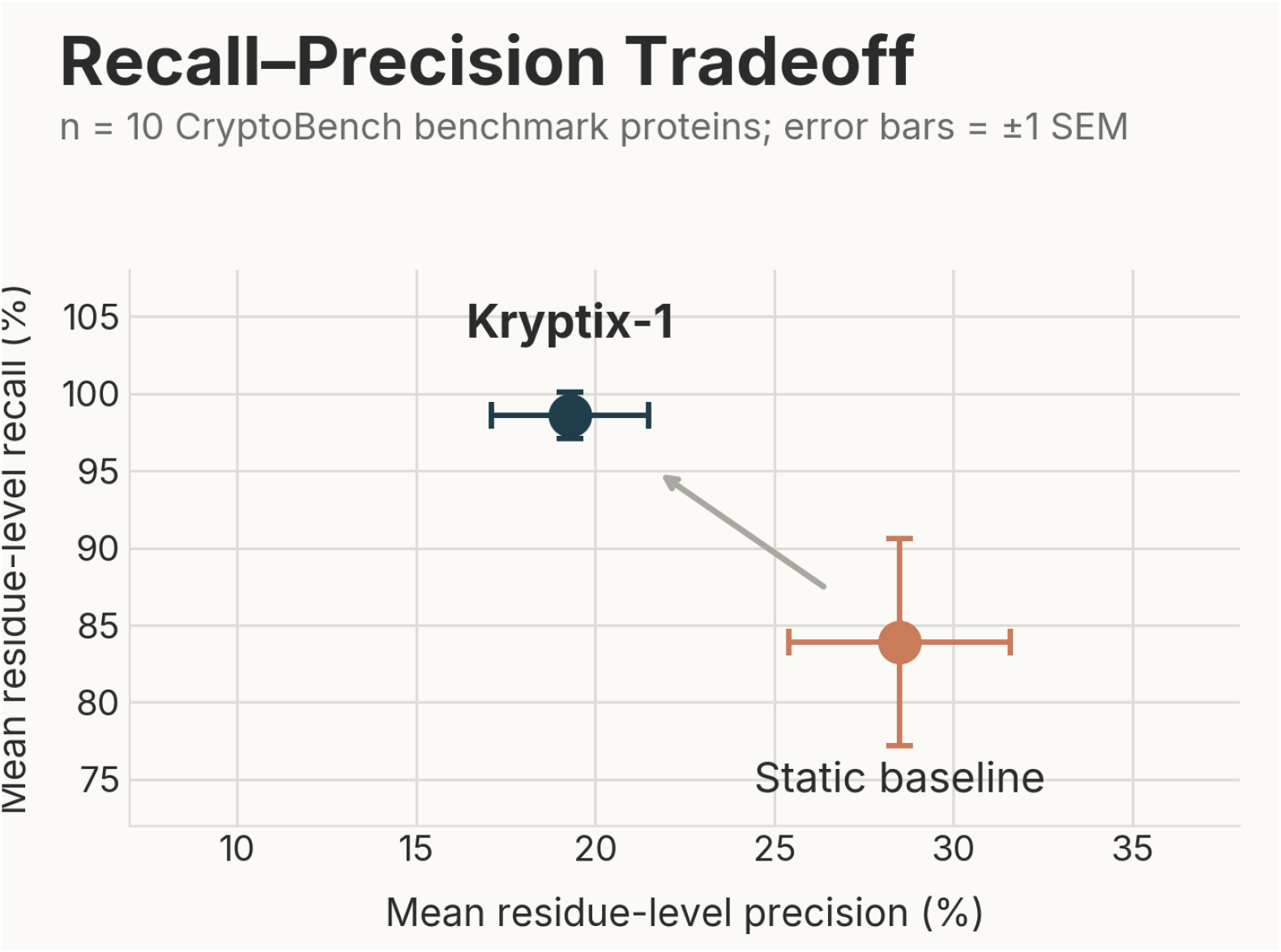
Recall–precision tradeoff, static baseline versus Kryptix-1, mean values across the 10-protein benchmark sample. Error bars show ±1 standard error of the mean (SEM); precision difference p = 0.005, recall difference p = 0.058 (paired t-test, n = 10).

## Discussion

### Kryptix-1 relative to prior published results

The comparison most directly relevant to Kryptix-1’s value proposition is against prior published studies reporting success rates on the same or closely comparable metrics, rather than against any single named tool. On the Jaccard overlap metric, arguably the most widely used residue-overlap success criterion in the pocket-prediction literature [17], independent comparative evaluations have consistently reported success rates clustering around 40–45% for the best-performing methods evaluated [17].

Kryptix-1’s 80% success rate on this same metric, evaluated on an independently curated benchmark dataset not used in its development, represents an approximately twofold improvement over this published ceiling. A similar pattern holds on the DCC metric at the strict 4 Å threshold. We regard this consistency across two independently defined metrics, rather than a strong result on a single favorable metric, as the more meaningful signal that the improvement reflects a genuine property of conformational-ensemble-based detection rather than an artifact of metric selection. We attribute this gap primarily to the fundamental limitation described in the

Introduction: methods that infer cryptic pocket propensity from a single static structure are attempting to detect a dynamical phenomenon from a single, and by definition non-cryptic-pocket-containing, snapshot [5, 15, 16]. Kryptix-1’s advantage is largest precisely where this limitation should matter most, recovering the correct general pocket location (Jaccard, DCC), rather than in fine-grained residue-level precision, consistent with the interpretation that ensemble sampling primarily helps a detector find pockets it would otherwise never see, rather than sharpen the boundary of pockets it would have found anyway.

### The recall–precision tradeoff, and why it does not undermine the core result

The Results section above shows lower mean precision for Kryptix-1 than for the static baseline, even as both success-rate metrics (Jaccard, DCC) and recall improve substantially. This is an expected consequence of ensemble-based detection: aggregating candidate signal across a larger number of sampled conformations increases the chance of recovering the true cryptic pocket, while also increasing the chance of flagging additional incorrect residues along the way. We report this tradeoff explicitly, rather than omitting it, because we believe transparency about where an approach does not dominate strengthens rather than weakens confidence in the results where it does.

We do not consider this tradeoff to undermine Kryptix-1’s practical value. The intended operational use of a cryptic-pocket screening tool is triage: surfacing a manageable shortlist of proteins and candidate sites worth committing downstream structural or experimental resources toward, not delivering final structural certainty on the first pass. For this purpose, the metrics that matter most are exactly the ones where Kryptix-1’s advantage is clearest and least ambiguous; whether the pocket is found at all, at approximately the correct location; rather than whether every individual flagged residue is correct. Given that inadequate target validation has been identified as a contributor to a substantial share of late-stage clinical failures [3, 4], we consider a tool that recovers substantially more true cryptic pockets, at moderate precision, to have a more favorable risk profile for a discovery pipeline than one that recovers fewer at marginally higher precision.

### Computational profile and scalability

The results reported here were generated using a single shared consumer-grade GPU. Because conformational sampling and pocket scoring are computed independently for each candidate protein; no protein’s analysis depends on the outcome of another’s; the pipeline is architecturally well suited to near-linear parallelization across additional compute capacity, consistent with reported scaling behavior for other GPU-accelerated stages of the drug discovery pipeline [20]. We expect that moving from a single shared consumer GPU to dedicated enterprise-scale infrastructure would reduce per-protein turnaround time by an order of magnitude or more; we state this as an architectural expectation grounded in the pipeline’s embarrassingly parallel structure across targets, not as a benchmarked result, since this scaling behavior has not yet been independently measured on such infrastructure.

### Limitations

This evaluation is based on a randomly drawn 10-protein sample from a single benchmark dataset, and we are explicit that it should be read as a pilot rather than a confirmatory study. It was not cherry-picked, and it includes proteins where Kryptix-1 shows a smaller advantage or a small reversal relative to the static baseline (Fig. 2), but a substantially larger validation set would be required to establish confidence intervals around the magnitude of improvement reported here, and to support subgroup analysis by protein size, fold class, or pocket burial depth. CryptoBench’s full published collection exceeds 1,100 structures [19] and represents a natural next validation step (see Conclusion and Outlook). We also note that the comparison to prior published success rates above, while using identical metric definitions and threshold conventions, was not conducted as a controlled head-to-head evaluation on an identical dataset; the reported published figures reflect the respective original studies’ own benchmark datasets and protocols, and should be read as context for the field’s current performance ceiling rather than a matched comparison.

## Methods

Kryptix-1’s specific algorithmic components; its conformational sampling strategy, model architecture, and consensus scoring logic; constitute proprietary intellectual property of Covenant Biosciences and are withheld from this report. At the architectural level, the pipeline operates in two stages: a conformational ensemble generation module, which samples plausible alternative structural states of an input protein beyond its static or crystallographic conformation, followed by a consensus pocket-scoring algorithm that aggregates candidate pocket signal across the generated ensemble into a final, ranked set of predicted cryptic sites, broadly in line with the consensus-scoring principle previously shown to improve reliability in the related virtual screening literature [18].

### Static-structure baseline

To isolate the specific contribution of conformational sampling from the scoring logic itself, all results are reported against a matched static-structure baseline: the identical consensus scoring algorithm applied to a single static input structure, with the ensemble generation stage disabled. Any performance difference between Kryptix-1 and the static baseline is therefore attributable specifically to conformational sampling, not to a difference in the underlying scoring methodology.

### Benchmark dataset

Ten proteins were randomly sampled from CryptoBench [19], a published cryptic-pocket reference dataset independently curated from paired apo/holo Protein Data Bank structures. Ground-truth pocket residues for each protein were taken directly from CryptoBench’s own annotated apo-structure pocket selections. No protein in the benchmark sample was used in the development, tuning, or calibration of Kryptix-1.

### Evaluation metrics

Three metrics were used, each adopted from established conventions in the independent pocket-prediction literature rather than defined for this report: Jaccard overlap (Irel success threshold) [17]; DCC at 4 Å (strict) and 12 Å (loose) thresholds [17]; and standard residue-level precision and recall. Binary success/failure outcomes (Jaccard, DCC) were compared between Kryptix-1 and the static baseline using an exact McNemar test on paired per-protein outcomes; continuous outcomes (precision, recall) were compared using a paired t-test and, given the small sample, corroborated with the non-parametric Wilcoxon signed-rank test. All tests are paired by protein (n = 10) and two-sided.

## Conclusion and Outlook

Kryptix-1 demonstrates that reintroducing conformational dynamics into cryptic pocket detection; rather than continuing to push single-structure predictors toward diminishing returns, or accepting the throughput ceiling imposed by exhaustive physics-based simulation; yields a substantial, metric-consistent improvement over both a matched static baseline and the published state of the art, while running on commodity hardware. We regard this as validation of the core architectural thesis rather than a finished product: the results reported here span ten proteins, one benchmark dataset, and a single consumer-grade GPU, and each of these constraints defines the immediate next phase of work.

Three priorities follow directly from the limitations noted above. First, extending this evaluation to CryptoBench’s full published collection will allow us to report the magnitude of Kryptix-1’s advantage with proper statistical confidence, and to characterize the specific structural features; pocket depth, fold class, apo/holo conformational displacement; that predict where ensemble sampling helps most. Second, we intend to pursue experimental validation of a subset of Kryptix-1’s novel predictions on internal protein targets, closing the loop between computational prediction and biophysical confirmation that this report, by design, does not address. Third, we plan to formally benchmark the scalability behavior described qualitatively in the Discussion on dedicated multi-GPU infrastructure, converting an architectural expectation into a measured result.

More broadly, we see this as early evidence that the precision-versus-throughput tradeoff long treated as fundamental to computational cryptic pocket discovery is an artifact of the specific methodological choices the field has made to date, and that closing it substantially, rather than only shifting where on the tradeoff curve a given tool sits, is achievable. We intend to pursue this direction as an internal discovery capability at Covenant Biosciences, and to publish updated, larger-scale results as the validation program described above matures.

## Data and Code Availability

The CryptoBench benchmark dataset used for evaluation is publicly available from its original publication [19]. The Kryptix-1 methodology, including its conformational sampling and scoring algorithms, is proprietary to Covenant Biosciences and is not disclosed or made available in connection with this report. Aggregate per-protein benchmark results underlying the figures and tables in this report are available from the corresponding author upon reasonable request.

## Author Contributions

All authors contributed to study design, implementation, analysis, and manuscript preparation.

## Acknowledgments

We thank the CryptoBench project for curating and publicly releasing the benchmark dataset used in this evaluation.

